# A bacteria-derived tail anchor localizes to peroxisomes in yeast and mammalian cells

**DOI:** 10.1101/437855

**Authors:** Güleycan Lutfullahoğlu-Bal, Ayşe Bengisu Seferoğlu, Abdurrahman Keskinb, Emel Akdoğan, Cory D. Dunna

**Author notes:** Current address: Department of Biological Sciences, Columbia University, New York, NY 10027, United States of America. Current address: Department of Microbiology and Molecular Genetics, University of California, Davis, Davis, CA 95616, United States of America. Corresponding author: Dr. Cory D. Dunn, Institute of Biotechnology, P.O. Box 56, 00014 University of Helsinki, Finland.

## Abstract

Prokaryotes can provide new genetic information to eukaryotes by horizontal gene transfer (HGT), and such transfers are likely to have been particularly consequential at the dawn of eukaryogenesis. Since eukaryotes are highly compartmentalized, it is worthwhile to consider the mechanisms by which newly transferred proteins might reach diverse organellar destinations. Toward this goal, we have focused our attention upon the behavior of bacteria-derived tail anchors (TAs) expressed in the eukaryote *Saccharomyces cerevisiae*. In this study, we report that a predicted membrane-associated domain of the *Escherichia coli* YgiM protein is specifically trafficked to peroxisomes in budding yeast, can be found at a pre-peroxisomal compartment (PPC) upon disruption of peroxisomal biogenesis, and can functionally replace an endogenous peroxisome-directed TA. Furthermore, the YgiM(TA) can also localize to peroxisomes in mammalian cells. Since the YgiM(TA) plays no endogenous role in peroxisomal function or assembly, this domain is likely to serve as an excellent tool toward illumination of the mechanisms by which TAs can travel to peroxisomes. Moreover, our findings emphasize the ease with which bacteria-derived sequences might target to organelles in eukaryotic cells following HGT, and we discuss the importance of flexible recognition of organelle targeting information during and after eukaryogenesis.

## INTRODUCTION

While prokaryotes can harbor compartments dedicated to specific functions and biochemical reactions ^1^, eukaryotes are commonly characterized by a higher level of compartmentalization by membranous structures. One of these organelles, the peroxisome, is bounded by a single membrane and is often a location of fatty acid oxidation in eukaryotic cells ^2,3^ Beyond fatty acid breakdown, peroxisomes play multiple roles among eukaryotes^4,5^, including sterol synthesis ^6^, synthesis of ether lipids ^7^, and even glycolysis ^8^. Soluble proteins are directed to the lumen, or matrix, of peroxisomes by a conserved import machinery commonly (but not exclusively) taking advantage of a carboxyl-terminal sequence called peroxisomal targeting sequence 1 (PTS1) ^9,10^. Membrane proteins are also targeted to peroxisomes, but mechanisms of peroxisomal membrane protein (PMP) biogenesis are not as well characterized as those processes that mediate import to the peroxisomal matrix ^11,12^. The evolutionary origin of peroxisomes is obscure, although some evidence suggests that the core machinery required for peroxisomal assembly is derived from the endoplasmic-reticulum-associated protein degradation (ERAD) machinery ^13,14^.

During and following eukaryogenesis, (proto-) nuclear genes were obtained by gene transfers from endosymbionts and from free-living prokaryotes, with some of these proteins subsequently targeted to organelles ^15–20^. Beyond more ‘ancient’ gene transfers, HGT from prokaryotes to eukaryotes and conversion of endosymbionts to organelles appears to continue at present day ^21–24^. Signals found within the polypeptide sequence of nucleus-encoded genes play a dominant role in targeting to eukaryotic organelles, and how prokaryote-derived proteins might acquire such sequences and become localized to eukaryotic organelles is a topic of intense inquiry. In a previous study ^25^ directed toward the principals of organelle targeting following HGT from bacteria, we focused our attention upon those proteins predicted to be anchored to membranes by a carboxyl-terminal hydrophobic stretch of amino acids, or a tail anchor (TA). Here, we describe the trafficking of one of these bacteria-derived TAs, retrieved from the YgiM protein of *E. coli.* We find that the YgiM tail anchor sequence [YgiM(TA)] localizes to peroxisomes in yeast and in human cells and can functionally replace an endogenous, peroxisomal TA in *S. cerevisiae.* In mutants in which peroxisomal biogenesis is impaired, the YgiM(TA) is localized to ER or to ER-derived peroxisomal compartments (PPCs), suggesting that this exogenous domain follows a trafficking pathway used by endogenously encoded peroxisomal TAs. Our work highlights the ability of eukaryotes to use prokaryotic information obtained by HGT to direct acquired proteins to distinct subcellular locations.

## RESULTS

### A domain encoded by the bacterial YgiM protein is targeted to the peroxisomes of yeast cells

During a previous appraisal of the ability of eukaryotic cells to utilize potential targeting information encoded by prokaryotes ^25^, we fused mCherry to the amino-terminus of TAs predicted to be encoded by the *E. coli* genome. These fluorescent fusion proteins were found at diverse locations within the cell, and we noted that mCherry fused to amino acids 173-206 of the uncharacterized YgiM protein, hereafter called the YgiM(TA), was found in a punctate pattern reminiscent of peroxisomes. The YgiM(TA) contains a predicted transmembrane helix followed by a positively charged lumenal tail (Fig. 1a). In order to determine whether the YgiM(TA) might indeed target to peroxisomes, we expressed mCherry-YgiM(TA) from the strong *ADH1* promoter together with superfolder green fluorescent protein (sfGFP) linked to the enhanced peroxisomal targeting signal 1 (ePTS1) ^26^. mCherry-YgiM(TA) co-localized with sfGFP-ePTS1, providing strong evidence of YgiM(TA) targeting to peroxisomes (Fig. 1 b). In contrast, mCherry-YgiM(TA) was not detectable at ER (Fig. 1c). Similarly, mCherry-YgiM(TA) was not detectable at mitochondria (Fig. 1 d), even upon deletion of Msp1p (Fig. S1), which extracts peroxisomal tail-anchored proteins mistargeted to mitochondria ^27,28^.

**Figure 1.**
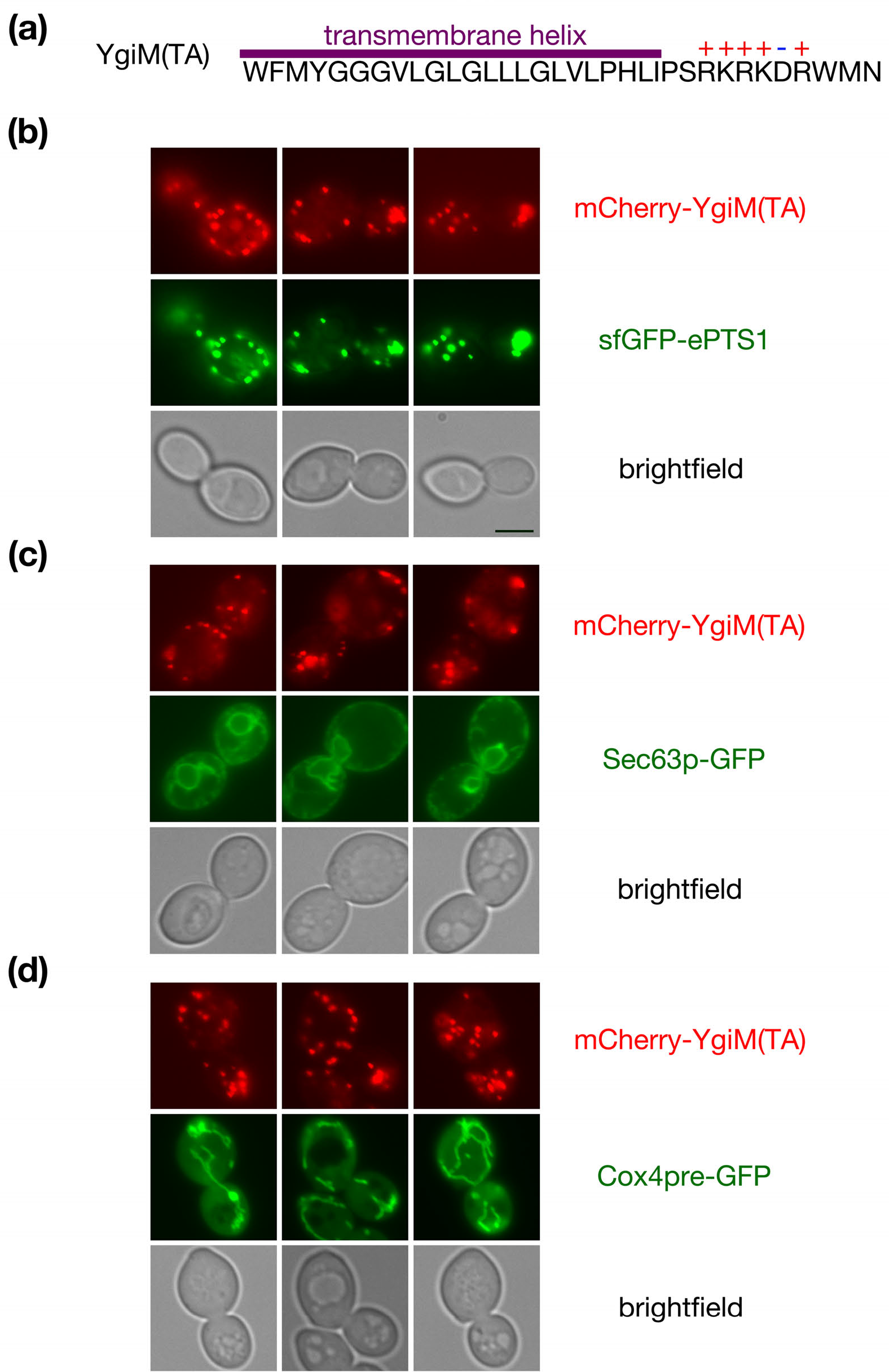
The predicted tail anchor of Escherichia coli YgiM localizes to peroxisomes in Saccharomyces cerevisiae. **(a)** The sequence of the YgiM(TA), as retrieved from UniProt^76^ record P0ADT8, is provided. The transmembrane helix is predicted using the TMHMM 2.0 server^77^ and charged residues are indicated, **(b)** The YgiM(TA) co-localizes with a protein targeted to peroxisomes. Strain BY4741, harboring plasmid b311 (sfGFP-ePTS1), was mated to strain BY4742, carrying plasmid b274 [mCherry-YgiM(TA)]. The resulting diploids were analyzed by fluorescence microscopy, **(c)** The YgiM(TA) does not co-localize with ER in wild-type cells. Strain BY4741, containing plasmid pJK59 (Sec63p-GFP) and strain BY4742, carrying plasmid b274 [mCherry-YgiM(TA)] were analyzed as in **(b). (d)** The YgiM(TA) does not co-localize with mitochondria. Strains BY4741 and BY4742, transformed with plasmids pHS1 (Cox4pre-GFP) and b274 [mCherry-YgiM(TA)], respectively, were mated and analyzed as in **(b).**

### The YgiM tail anchor can functionally replace an endogenous, peroxisome-directed tail anchor

Pex15p, which participates in the import of yeast proteins to the peroxisomal matrix ^9,10,29^, is the only *S. cerevisiae* protein thought to be directed specifically to peroxisomes by a TA ^30^. A lack of Pex15p at peroxisomes leads to defective peroxisomal biogenesis and cytosolic accumulation of PTS1-directed proteins ^31^. Previous studies have demonstrated that Pex15p is functional when its TA is replaced by that of the mammalian PEX26 protein ^32^, suggesting that other peroxisome-inserted TAs might also support Pex15p activity. Therefore, we tested whether the YgiM(TA) might target the Pex15p cytosolic domain to peroxisomes and permit Pex15p-driven peroxisomal protein import.

As expected, expression of an untethered Pex15p cytosolic domain (amino acids 1-331) ^32^ under control of the native *PEX15* promoter in cells lacking a chromosomal copy of *PEX15* did not allow peroxisomal import of sfGFP-ePTS1 (Fig. 2a and 2e), while re-attachment of the Pex15(TA) to the Pex15p cytosolic domain permitted efficient import of the same substrate to the peroxisomal matrix (Fig. 2b and 2e). Demonstrating that the bacterial YgiM(TA) can provide functionality in S. *cerevisiae*, Pex15(1-331)-YgiM(TA) allowed peroxisomal import of sfGFP-ePTS1, although rescue of the *pex15Δ* phenotype was not absolute (Fig. 2c and 2e). Not all bacteria-derived TAs can support Pex15p function: Pex15(1-331) fused to the *E. coli* YqjD(TA), which was previously demonstrated ^25^ to target predominantly to mitochondria and, to a lesser extent, the ER, failed to allow *pex15Δ* rescue (Fig. 2d and 2e). Though a portion of the S. *cerevisiae* Fis1 p is associated with peroxisomes ^33^, we found no evidence that the Fis1 p(TA) can support Pex15p function (Fig. S2).

**Figure 2.**
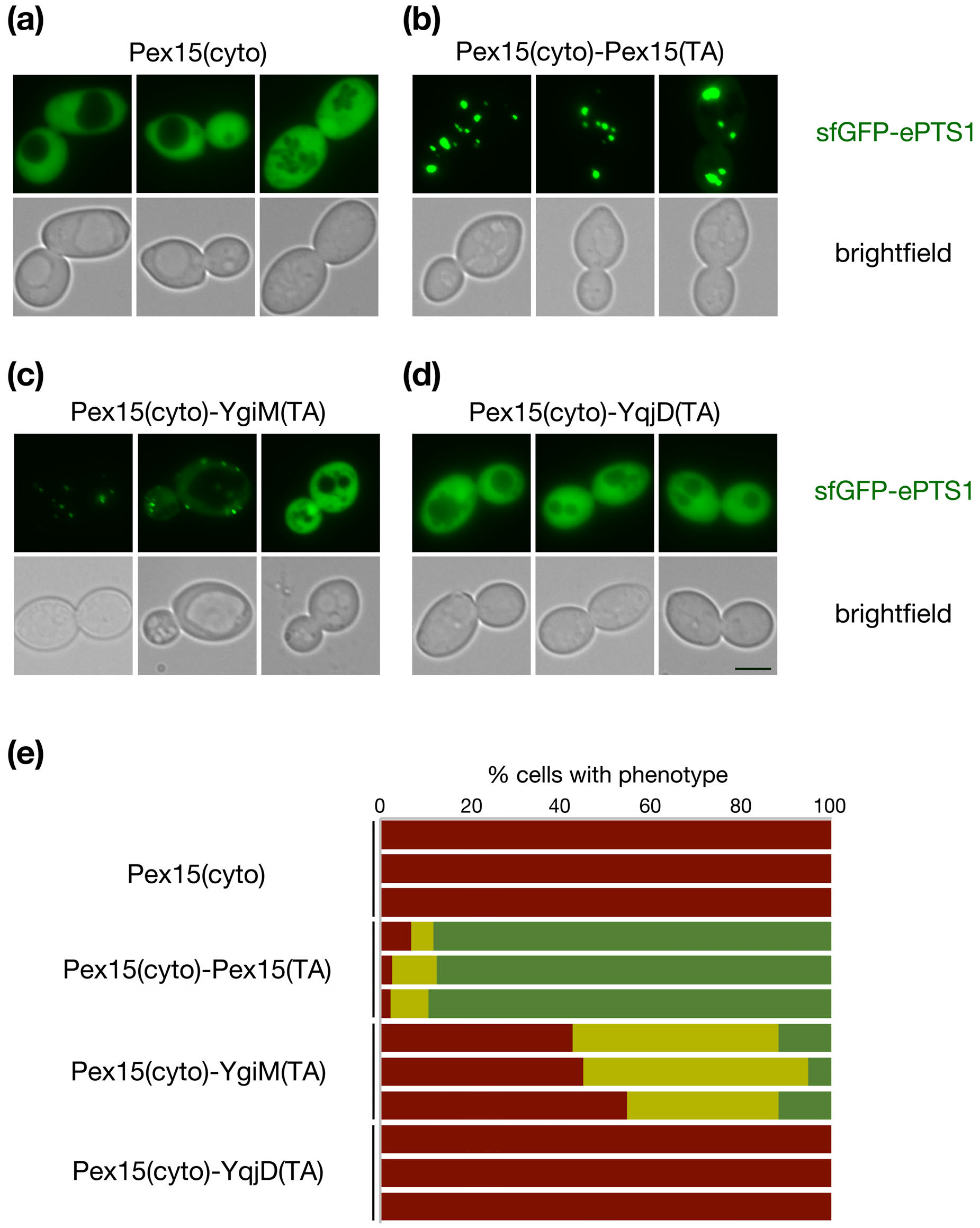
The YgiM(TA) can functionally replace the tail anchor of Pex15p. *pex15Δ/pex15Δ* strain CDD1182, containing a counter-selectable plasmid expressing the Pex15p cytosolic domain (cyto) fused to its own TA (b354) was transformed with plasmids expressing **(a)** Pex15(cyto) lacking a TA (b341) **(b)** Pex15(cyto)-Pex15(TA) (b326) **(c)** Pex15(cyto) fused to the YgiM(TA) (b329) or **(d)** Pex15(cyto) fused to the YqjD(TA) (b330). Plasmid b354 was then removed by counter-selection with CHX, and Pex15p function was assessed by sfGFP-ePTS1 localization to peroxisomal puncta. **(e)** Reports the quantification, blinded to genotype during analysis, of three independent experiments. Red represents cells with diffuse signal in the nucleus and cytosol but no puncta, yellow represents cells with both diffuse and punctate signal, and green represents cells in which only punctate, peroxisomal signal representing peroxisomes could be detected (n>200 cells per sample in each experimental replicate).

### The YgiM tail anchor resides within a pre-peroxisomal compartment upon disruption of peroxisomal biogenesis

In yeast, many integral peroxisomal membrane proteins (PMPs) are inserted at the ER before subsequent trafficking to peroxisomes ^12^. The tail-anchored Pex15 protein is also thought to begin its journey to the peroxisome within the ER ^30,34^, and upon disruption of PMP trafficking, accumulates in ER-derived pre-peroxisomal compartments (PPCs) ^35^ marked by Pex14p ^36–38^, a component contributing to formation of the mature peroxisomal import pore ^39,40^.

To visualize Pex14p-marked PPCs, we tagged endogenous Pex14p with sfGFP. The Pex14p-sfGFP fusion protein was easily detectable, could promote peroxisomal protein import (Fig. S3a), and continued to be localized, as previously reported, in puncta representing PPCs upon disruption of peroxisomal biogenesis by removal of Pex3p or Pex19p (Fig.S3b and S3c). Some mCherry-Pex15(TA) was found to co-localize with Pex14p at peroxisomes of WT cells (Fig. S4a), although a notable fraction of mCherry-Pex15(TA) is also mistargeted to mitochondria (Fig. S4d). Consistent with a previous report examining the trafficking of full-length Pex15p ^35^, we found that mCherry-Pex15(TA) could be co-localized with Pex14-sfGFP upon disruption of PMP trafficking by removal of Pex3p (Fig. S4b) or Pex19p (Fig. S4c).

If YgiM(TA) is, like endogenous PMPs, initially targeted to ER, it might similarly be localized to PPCs upon disruption of PMP trafficking. mCherry-YgiM(TA) co-localized with Pex14p-sfGFP at mature peroxisomes in wild-type cells (Fig. 3a), and indeed, mCherry-YgiM(TA) continued to co-localize with Pex14p-sfGFP in *pex3Δ* (Fig. 3b) or *pex19Δ* (Fig. 3c) cells. Our findings are consistent with trafficking of the YgiM(TA) to the ER, then to PPCs, before subsequent movement to peroxisomes, and our results suggest consonance between cellular pathways handing the endogenous Pex15(TA) and the bacterial, exogenous YgiM(TA).

**Figure 3.**
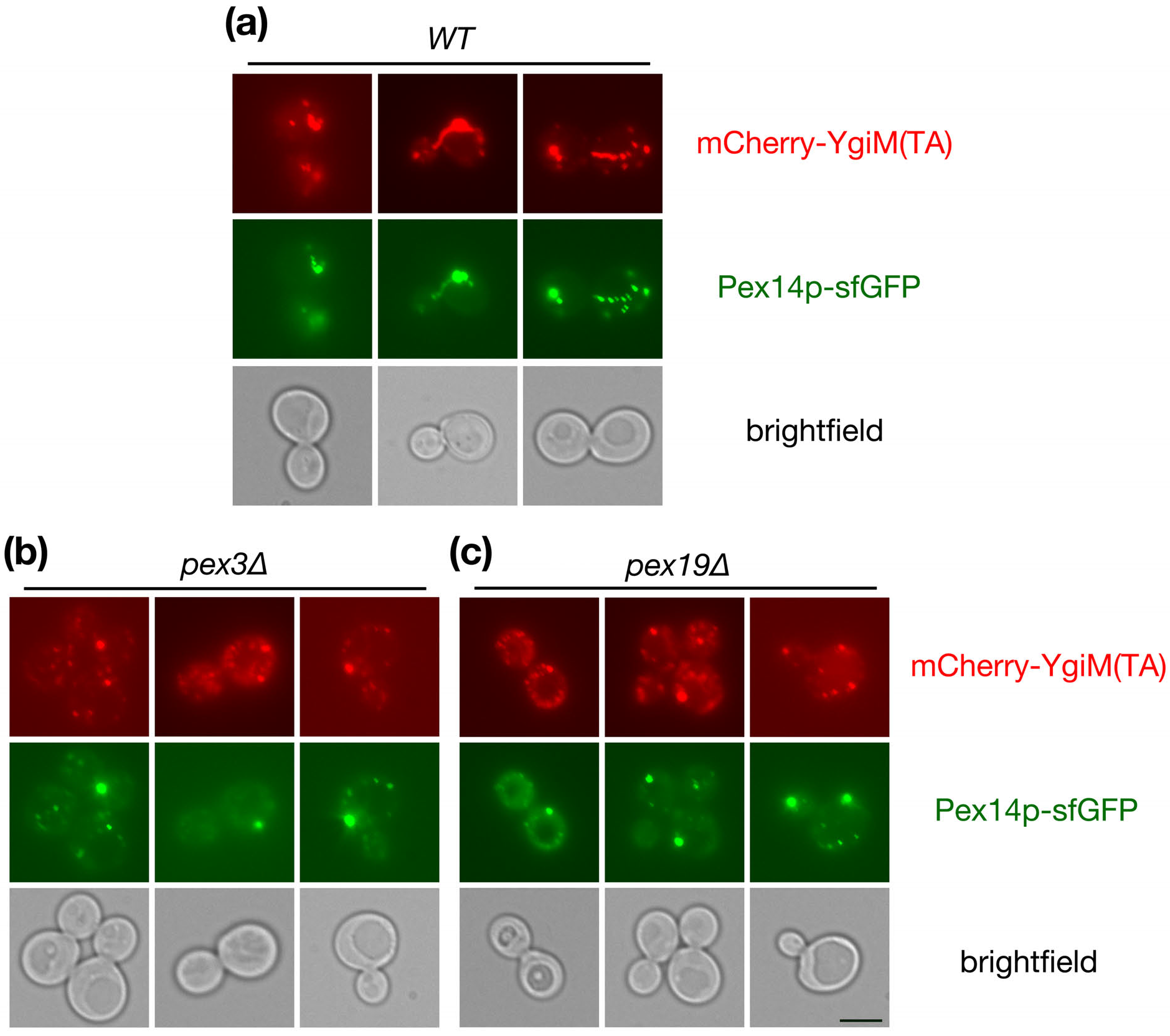
The YgiM(TA) localizes to PPCs containing Pex14p upon blockade of PMP transit to peroxisomes. **(a)** WT (CDD1200) **(b)** *pex3Δ* (CDD1201) or **(c)** *pex19Δ* (CDD1202) cells expressing mCherry-YgiM(TA) from plasmid b274 and Pex14p-sfGFP from the native *PEX14* locus were examined by fluorescence microscopy.

### The ER-localized Spf1 protein contributes to trafficking of the YgiM tail anchor to peroxisomes

Spf1p, an ER-localized protein involved in manganese transport ^41^, plays a role in peroxisomal biogenesis ^42,43^, and the localization of at least two proteins capable of trafficking from ER to peroxisomes, Pex3p and Ant1p ^44–47^, is altered by Spf1p removal ^48^. Consequently, we investigated whether trafficking of YgiM(TA), like endogenously encoded PMPs, might also be affected by Spf1 p deletion. Indeed, mCherry-YgiM(TA) was significantly redistributed to ER in *spf1Δ* cells (Fig. 4a), demonstrating a potential role for Spf1p in the targeting of peroxisome-directed TAs and consistent with YgiM(TA) trafficking through the ER. Interestingly, mCherry-YgiM(TA) was also found at mature peroxisomes marked by sfGFP-ePTS1 in *spf1Δ* cells, demonstrating that Spf1p removal does not completely abolish TA trafficking. We also note that Spf1p is not apparently required for the generation of PPCs containing Pex14p, since Pex14p-sfGFP puncta are easily visualized in *spf1Δ* cells, including within cells also deleted of Pex3p or Pex19p (Lutfullahoğlu-Bal G, unpublished data).

**Figure 4.**
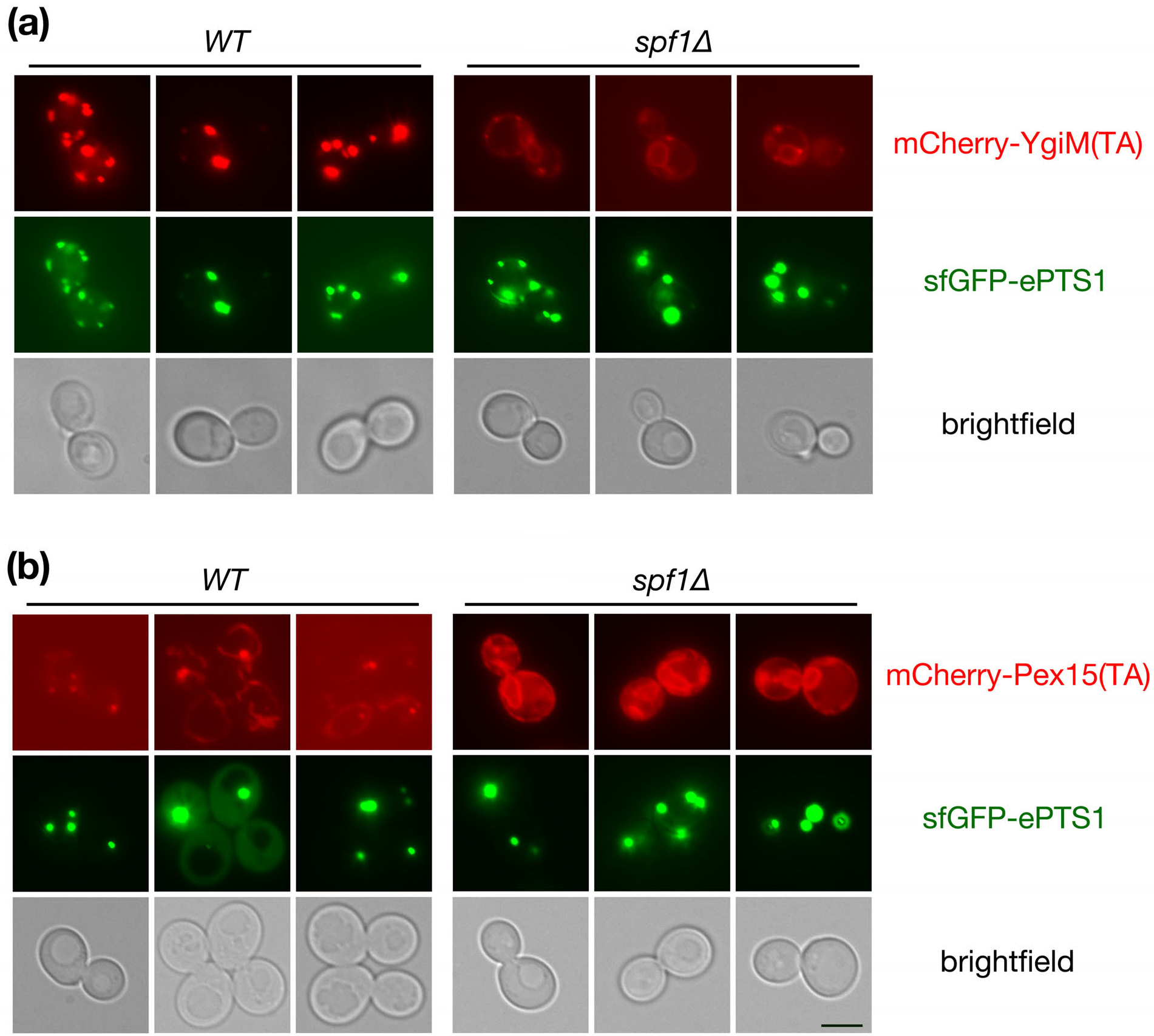
Peroxisome-directed tail anchors are mislocalized upon deletion of the Spf1 protein. **(a)** mCherry-YgiM(TA), expressed from plasmid b274, or **(b)** mCherry-Pex15(TA) expressed from plasmid b365 was expressed in *WT* (BY4742) or *spflΔ* (CDD949) cells. Cells also expressed sfGFP-ePTS1 from plasmid b311 in order to determine the location of peroxisomes.

We tested whether the endogenously encoded Pex15(TA) would, like the YgiM(TA), be redistributed across the ER upon deletion of Spf1 p. Indeed, like mCherry-YgiM(TA), mCherry-Pex15(TA) was localized abundantly to ER in *spf1Δ* cells but not in WT cells (Fig. 4b), again indicating congruence in trafficking mechanisms used by the YgiM(TA) and the Pex15(TA).

### Expression of the YgiM(TA) does not disturb peroxisomal biogenesis

Since overexpression of full-length Pex15p perturbs peroxisomal biogenesis ^30^, we asked whether expression of only the TA domains of YgiM or Pex15p, also driven by the strong *ADH1* promoter, would have a detrimental effect on peroxisome assembly. Toward this goal, the behavior of sfGFP-ePTS1 was assessed in cells expressing mCherry-YgiM(TA) or mCherry-Pex15(TA). Peroxisomal biogenesis was indeed disrupted by mCherry-Pex15(TA) overexpression (Fig. 5), with an average of 8% of cells lacking discernable peroxisomes across three independent experiments. Moreover partial nucleocytoplasmic accumulation of sfGFP-ePTS1 was visible in nearly twice as many cells expressing mCherry-Pex15(TA) as those expressing empty vector. However, mCherry-YgiM(TA) expression had no effect on sfGFP-ePTS1 localization when compared to cells harboring empty vector; distinct peroxisomes could be visualized in all cells. Therefore, overexpression of the YgiM(TA), unlike overexpression of an endogenous peroxisome-directed TA, does not appear to disrupt peroxisomal biogenesis.

**Figure 5.**
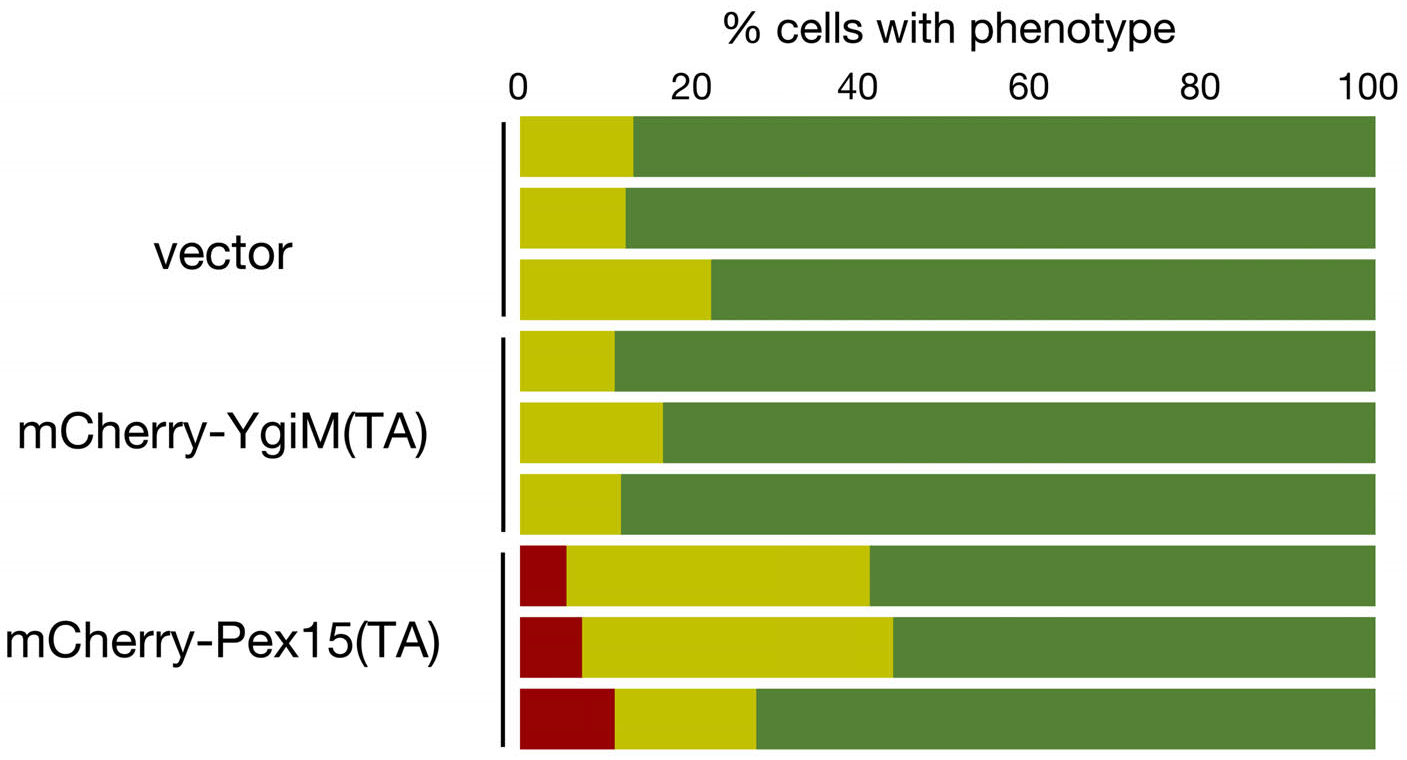
Overexpressed Pex15(TA), but not overexpressed YgiM(TA), perturbs peroxisomal biogenesis. *WT* cells (BY4742) expressing sfGFP-ePTS1 from plasmid b311 along with mCherry-YgiM(TA) from plasmid b274, mCherry-Pex15(TA) from plasmid b365, or empty vector pRS315 were examined as in Fig. 2e.

### The YgiM(TA) is localized to the peroxisomes of mammalian cells

Finally, we investigated whether YgiM(TA) might target to peroxisomes in mammalian cells, since the mechanism by which tail-anchored proteins are delivered to peroxisomes may differ between yeast and mammals ^9,12,49^. Upon transient transfection of a construct in which enhanced green fluorescent protein (EGFP) is fused to the YgiM(TA), punctate structures suggesting peroxisomal localization were visualized in HEK293T cells (Fig. 6a). These structures were confirmed to be peroxisomes upon co-localization with catalase, a marker of mature peroxisomes. As in yeast, EGFP-YgiM(TA) was not trafficked to mitochondria, as revealed by a lack of co-localization between EGFP-YgiM(TA) and the mitochondrial T0M20 protein (Fig. 6b).

**Figure 6.**
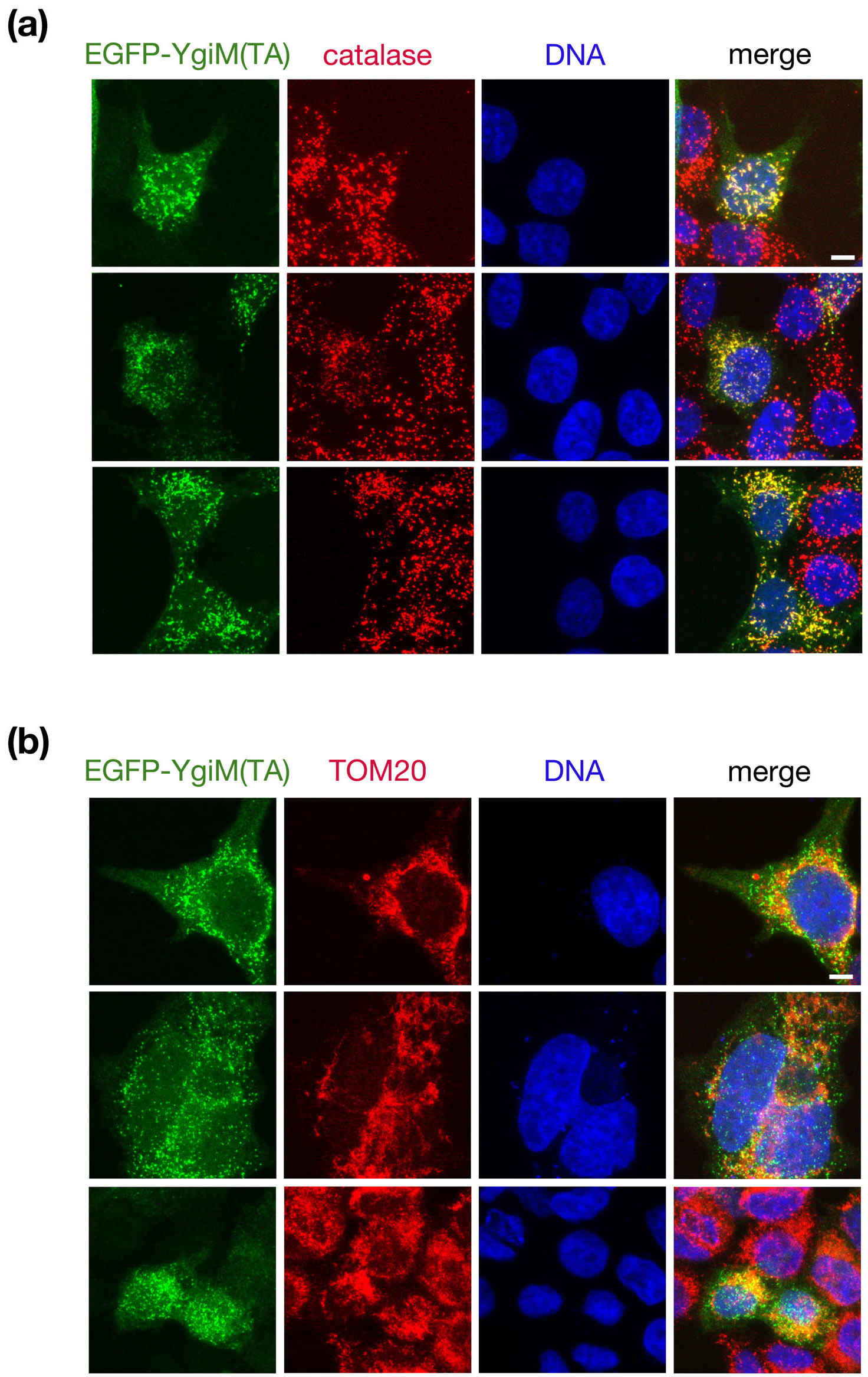
The YgiM(TA) localizes to peroxisomes in mammalian cells. Plasmid b374, expressing EGFP-YgiM(TA), was transiently transfected into HEK293T cells, and cells were processed for immunofluorescence. Anti-GFP antibodies were used to detect EGFP-YgiM(TA), and DAPI was used to stain cellular DNA. Anti-catalase antibodies were used to label mature peroxisomes **(a)** or anti-TOM2Q antibodies were used to label mitochondria **(b).**

## DISCUSSION

### What features of YgiM(TA) allow targeting to peroxisomes?

The *E. coli* YgiM(TA) was detected at peroxisomes, with no microscopic or functional evidence of mitochondrial localization in yeast or in human cells. Conversely, TAs found within two other proteins encoded by the same organism, YqjD and ElaB, targeted to mitochondria and ER, with no evidence of peroxisomal localization ^25^ [and this study]. Other tail-anchored proteins can target to both mitochondria and peroxisomes. For example, human Fis1 ^50,51^, yeast Fis1p ^33^, and human Mff can be localized to both organelles ^52^. The parameters that allow TAs to discriminate between peroxisomes, mitochondria, and other organelles are not understood, but may be related to the hydrophobicity of the membrane associated domain, along with the number and specific location of charges within the TA ^53,54^. When considering the recent development of a classifier for peroxisome-directed mammalian TAs ^54^, the GRAVY hydrophobicity score (1.7) ^55^ of the YgiM transmembrane domain, denoting more limited hydrophobicity, together with the net charge (+4.1) of the proposed lumenal tail at neutral pH (Protein Calculator v3.4, http://protcalc.sourceforge.net/) do, in fact, predict peroxisomal localization of the YgiM(TA). We note that the biogenesis of YgiM in bacteria has not been investigated, and indeed full-length YgiM contains a predicted signal sequence at its amino-terminus ^56^, indicating co-translational insertion and suggesting that any predicted TA would not drive initial membrane targeting in *E. coli.*

### Heterologous expression of YgiM(TA) may reveal mechanisms of tail-anchor trafficking to peroxisomes

Only two peroxisome-directed proteins harboring TAs are encoded by S. *cerevisiae:* Pex15p and Fis1p. Both associate with other peroxisomal proteins: Pex15p is found in a complex with Pex3p ^57^, and Fis1p cooperates with Pex11p ^58^. Based on these findings, one might propose a scenario in which both tail-anchored polypeptides obtain their final peroxisomal location solely through their functional assembly with other proteins not harboring a TA, and that no pathway with a specific role in directing tail-anchored client proteins to peroxisomes exists in budding yeast. However, the YgiM(TA), separated from eukaryotes by billions of years of evolutionary distance, appears to localize specifically to peroxisomes in both yeast and human cells. This exogenously expressed domain would not bind to any endogenous interaction partners in order to carry out a cellular function, yet makes its way to peroxisomes nonetheless, supporting the presence of a more generalized mechanism that allows trafficking of tail-anchored proteins to peroxisomes in *S. cerevisiae*.

Importantly, expression of native proteins at incorrect stoichiometry can perturb cellular functions that may be under investigation ^59,60^, as illustrated by disruption of peroxisomal biogenesis upon overexpression of full-length Pex15p ^30^. In this study, we found that the TA of Pex15p could also disrupt peroxisomal protein import, attenuating its value as an experimental substrate for studies of TA targeting in yeast. In contrast, YgiM(TA) expression did not affect peroxisomal assembly. Moreover, the YgiM(TA) is also relatively short when compared to several other peroxisome-directed TAs, providing the opportunity for facile mutational analyses of YgiM(TA). Therefore, we suggest that heterologously expressed YgiM(TA) is likely to be a preferred substrate for further exploration of the mechanisms by which TAs reach peroxisomes.

### Why is protein targeting to eukaryotic organelles so permissive?

In this study, we have demonstrated that a predicted membrane insertion sequence obtained from a prokaryote can be directed to the peroxisomes of eukaryotic cells. Our findings expand upon earlier studies in which protein sequence derived from prokaryotes could traffic to eukaryotic organelles, such as ER and mitochondria ^25,61–65^. We propose that the ability to direct prokaryotic protein sequences to eukaryotic organelles, even though these regions were not previously selected for targeting prowess in eukaryotes, might have been of general benefit to eukaryotes over evolutionary time. Specifically, failure to allow degeneracy among organelle targeting sequences ^66^ would potentially have limited the utility of genes acquired from the proto-mitochondrial endosymbiont or from neighboring microorganisms occurring at or near the dawn of eukaryogenesis, potentially slowing or forbidding the emergence of the eukaryotic cell ^67^. In addition, the ability of eukaryotes to take advantage of genes acquired by HGT at present day ^23,68,69^ could similarly be hampered by a strict sequence requirement, rather than lax structural requirements, for recognition of organelle-targeting signals contained within polypeptides. Additionally, sequestration at an organelle might avoid detrimental effects of aggregation or chaperone sequestration, and thereby avoid selection against an otherwise advantageous gene transfer, when taking into consideration hydrophobic regions like the TA examined in this study.

Encompassing the specific use of HGT-acquired genetic information would be a more general need for eukaryotic cells to harbor permissive organelle translocation machineries that allow recognition of degenerate import signals. Given that organelles are maintained in order to compartmentalize biochemical pathways and other cellular activities, it follows that multiple polypeptides will often act together as a module within a given organelle. Strict sequence requirements for organelle import would make it highly improbable that multiple proteins that once cooperated together at one cellular location, such as the cytosol, could later find themselves simultaneously localized together in a different cellular compartment. Conversely, more relaxed structural determinants of organelle-targeting regions of a protein that might be recognized by permissive substrate receptors, such as hydrophobicity and charge, would allow proteins already encoded by a cell to sample novel compartments. Eventually, as previously proposed by Martin ^70^, organelle sampling by polypeptides, followed by further mutational tinkering of the organelle targeting sequences, could lead to increased fitness through the localization of an entire cellular pathway to a new location. Moreover, genes can evolve *de novo* ^71^, and the ability of newly generated polypeptides to test different organelle environments may also contribute to fitness. Ultimately, then, the question of how the protein translocation machineries of organelles recognize targeting information of client proteins, obtained by HGT or as the outcome of other genetic processes, becomes a question of ‘evolvability,’ or the advantageous capacity of a pedigree of organisms to more easily sample genotypic and phenotypic space ^72^.

## METHODOLOGY

### Yeast strains, plasmids, and culture conditions

Culture conditions are as described in ^73^. All experiments with S. *cerevisiae* have been performed at 30°C. Plasmids, strains, and oligonucleotides used in this study can be located in Tables S1a, S1b, and S1c, respectively.

### Assessment of Pex15 functionality

Diploid strain CDD1182, deleted of chromosomal *PEX15*, expressing peroxisome-targeted sfGFP from plasmid b311, and carrying a fully-functional fusion between the cytosolic and TA domains of Pex15p from plasmid b354 driven by the *PEX15* promoter, was transformed with plasmids expressing variants of Pex15p in which the cytosolic domain was fused to test TAs by a linker region consisting of Fis1p amino acids 119-128, a stretch of amino acids not necessary or sufficient for organelle targeting ^74,75^. Strains were cultured overnight in supplemented minimal medium (SMM) lacking uracil and histidine (-Ura-His). Cells were then transferred to SMM-Ura-His containing 3 mg/L cycloheximide (CHX) and cultured overnight before fluorescence microscopy in the logarithmic phase of proliferation. Counter-selection of plasmid b354 was confirmed by lack of proliferation on medium lacking tryptophan.

### Mammalian cell culture and transfection

Cells were maintained at 37°C and 5% CO_2_ and cultured in Dulbecco’s Modified Eagle’s Medium (DMEM) supplemented with 10% fetal bovine serum, 2 mM L-glutamine, 100 U/ml penicillin/streptomycin, and 50 μg/ml uridine.

HEK293T cells were plated overnight before transfection in 500 μl of complete growth medium at a cell density of 1 × 10^5^ cells/ml in a 24-well plate containing glass coverslips. Transfection was performed using TransIT-2020 (Mirus Bio) reagent, and transfection mixture contained: 250 ng of plasmid b374, 50 μl of cell culture medium, and 1 μl of transfection reagent. The mixture was incubated at room temperature for 20 min, and transfection mixture was added drop-wise to the cells. Cells were fixed for immunofluorescence analysis 24 hrs after transfection.

### Microscopy

Microscopy on yeast cells was performed using logarithmic phase cultures, as in ^25^. mCherry fusions are driven by the *ADH1* promoter and contain Fis1p amino acids 119-128 linking mCherry to each carboxyl-terminal organelle targeting sequence.

To carry out indirect immunofluorescence experiments, transfected cells were fixed using 4% paraformaldehyde in phosphate-buffered saline (PBS), pH 7.4 for 10 min at room temperature. Cells were washed three times with PBS for 5 min, then blocked for 1 hr using PBS containing 0.3% Triton X and 1% bovine serum albumin. Cells were then incubated overnight in primary antibodies (listed in Table S1d) diluted in blocking solution at 4°C. Cells were washed 3x with PBS, 5 min each wash. Cells were incubated with secondary antibodies in the blocking solution for 1 h in the dark, and after secondary antibody incubation, 4′,6-diamidino-2-phenylindole (DAPI) was added to a final concentration of 1 μg/ml for 10 min. Cells were washed 3x with PBS, and coverslips were mounted using 80% glycerol prepared in 20 mM Tris-HCl pH 8.0. Coverslips were sealed and stored at 4°C before microscopy. Imaging was performed using a Zeiss LSM700 Axio Imager.M2 confocal microscope equipped with an LCI Plan-Neofluar 63x/1.30 Imm Corr objective using emission/excitation detection wavelengths of 405nm/435nm, 488nm/518nm, or 555nm/585nm. Scale bars provided with yeast and mammalian cell microscopy images correspond to 5 μm.

## ACKNOWLEDGEMENTS

We thank Pekka Katajisto (University of Helsinki) for anti-catalase antibodies, and we thank Gülayşe ince Dunn (University of Helsinki and Koç University) for HEK293T cells. In addition, we thank Maya Schuldiner, Ville Paavilainen, Ani Akpinar, and Gülayşe ince Dunn for helpful comments on this manuscript. This work was supported by a European Research Council Starting Grant (637649-RevMito) to C.D.D., by a Turkish Academy of Sciences Outstanding Young Scientist Award (TÜBA-GEBiP) to C.D.D., by an EMBO Installation Grant (2138) to C.D.D., and by KoKoç University. These funding bodies had no role in the design of the study, data collection, data analysis, data interpretation, or manuscript preparation.

## AUTHOR CONTRIBUTIONS

CDD designed the study, wrote the manuscript, and performed experiments. GLB and ABS performed experiments and generated reagents. AK and EA generated reagents. All authors read and approved the final manuscript.

## ABBREVIATIONS

CHX: Cycloheximide
DAPI: 4′,6-diamidino-2-phenylindole
DMEM: Dulbecco’s Modified Eagle’s Medium
EGFP: Enhanced Green Fluorescent Protein
ePTS1: Enhanced Peroxisomal Targeting Signal 1
ER: Endoplasmic Reticulum
ERAD: Endoplasmic-Reticulum-Associated Protein Degradation
HGT: Horizontal Gene Transfer
PBS: Phosphate-Buffered Saline
PMP: Peroxisomal Membrane Protein
PPC: Pre-Peroxisomal Compartment
PTS1: Peroxisomal Targeting Signal 1
sfGFP: Superfolder Green Fluorescent Protein
SMM: Supplemented Minimal Medium
TA: Tail Anchor

---

**Figure S1.**
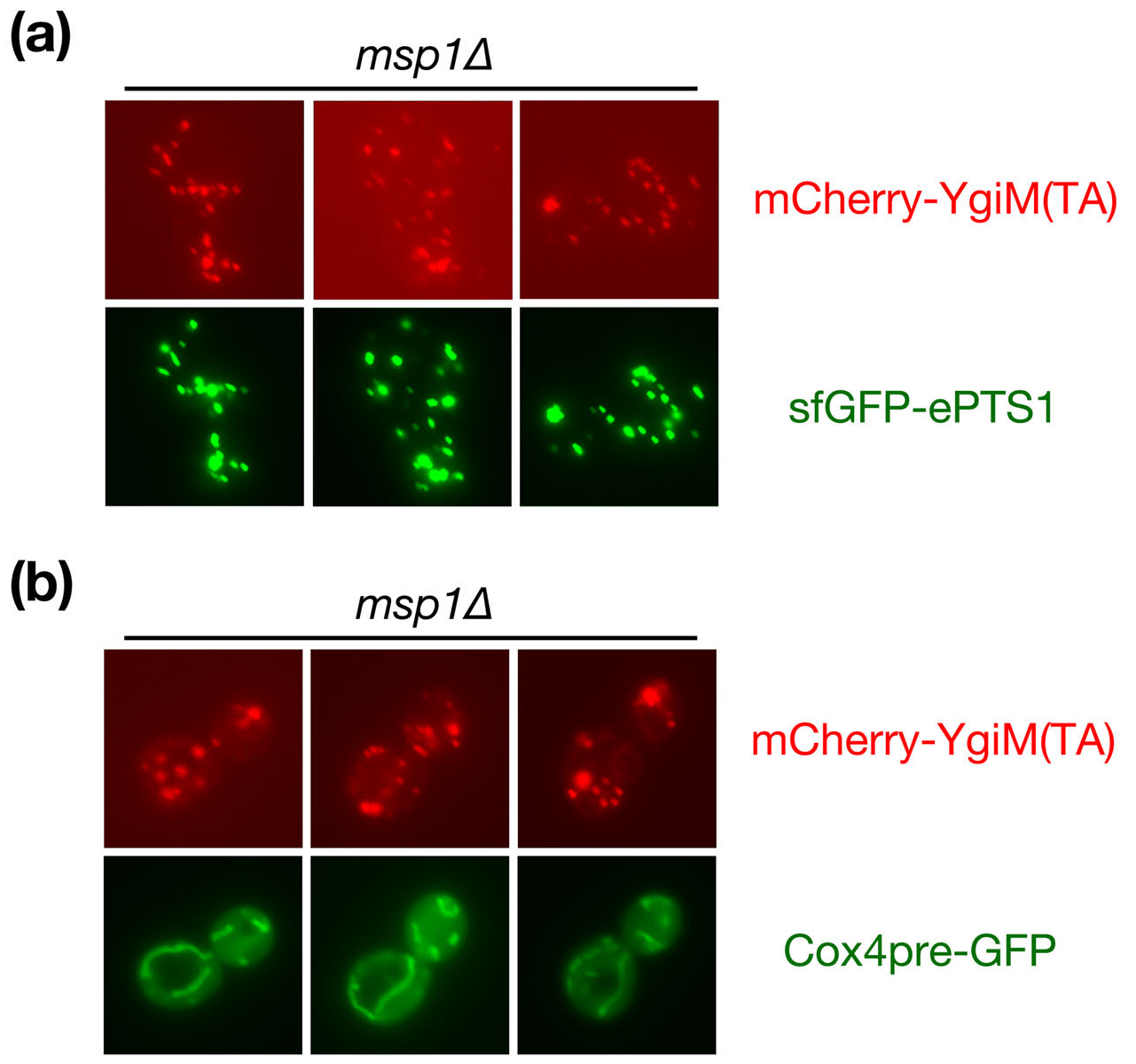
The YgiM(TA) does not accumulate at mitochondria upon deletion of Msplp. mCherry-YgiM(TA), encoded by plasmid b274 was expressed in *msplΔ* (CDD1154) cells, **(a)** Peroxisomes were labelled with sfGFP-ePTS1 expressed from plasmid b311, or **(b)** mitochondria were labelled by Cox4pre-GFP expressed from plasmid pHS1. Cells were examined by fluorescence microscopy.

**Figure S2.**
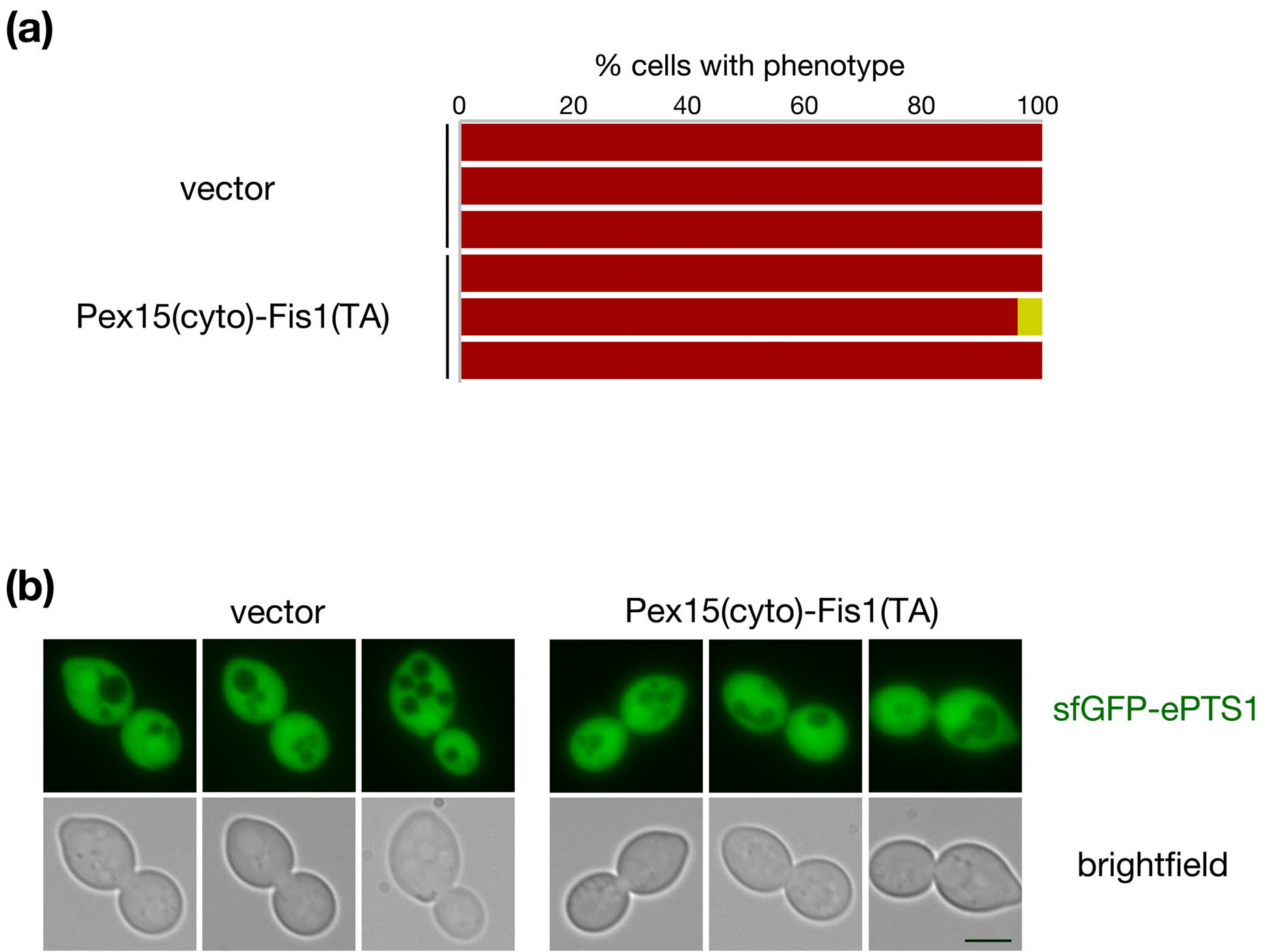
The Pex15 cytosolic domain fused to the Fislp TA is not functional. **(a)** *pex15Δ/pex15Δ* strain CDD1182 was transformed with plasmids expressing the Pex15p cytosolic domain (cyto) fused to the Fis1 p(TA) (b327) or with empty vector pRS316, and plasmid b354 was removed by counter-selection. sfGFP-ePTS1 expressed from plasmid b311 was examined as in Fig. 2e. Representative images are provided in **(b).**

**Figure S3.**
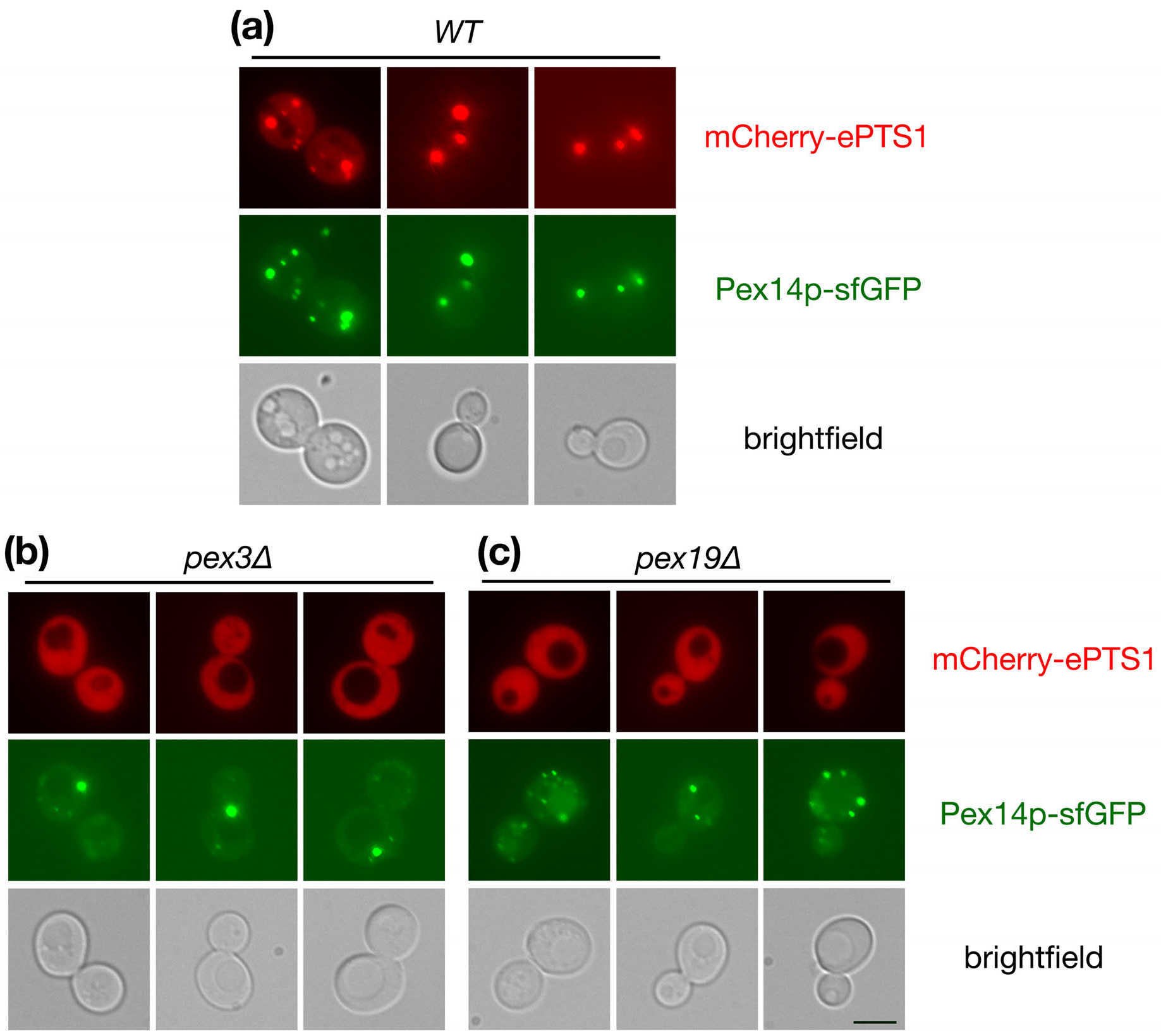
The Pex14p-sfGFP fusion protein is functional and is detectable in mutants in which PMP trafficking is blocked. *WT* (CDD1200), pex3A (CDD1201), and *pex19Δ* (CDD1202) strains, all harboring a chromosomal *PEX14-sfGFP* allele, were transformed with plasmid b364 expressing mCherry-ePTS1.

**Figure S4.**
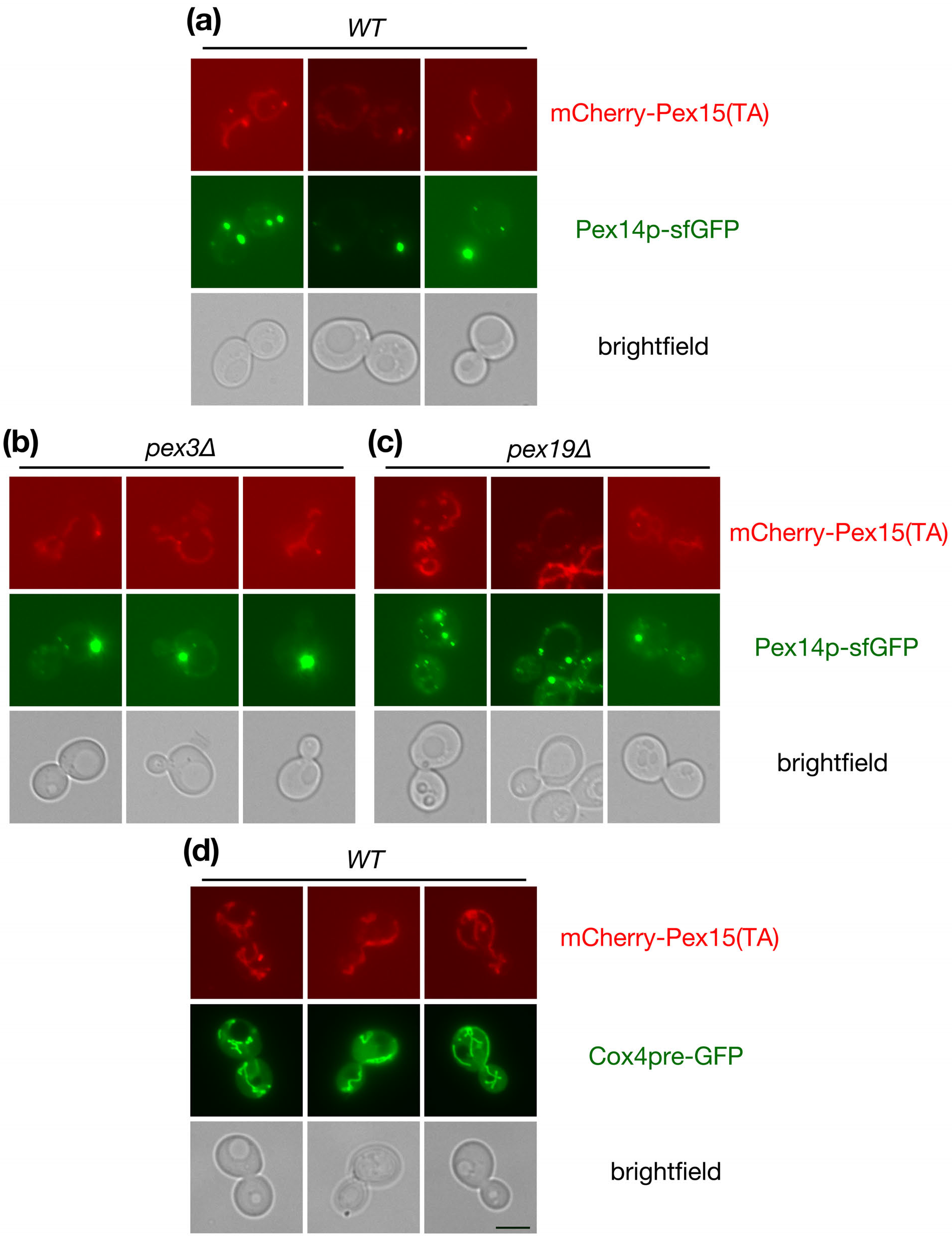
The Pex15(TA) can be found at Pex14p-positive PPCs upon disruption of PMP trafficking to peroxisomes. **(a)** *WT* (CDD1200) **(b)** *pex3Δ* (CDD1201) or **(c)** *pex19Δ* (CDD1202) cells expressing mCherry-Pex15(TA) from plasmid b365 were examined by fluorescence microscopy as in Fig. 3. **(d)** The Pex15(TA) is partially mislocalized to mitochondria in wild-type cells. Strain BY4741 was transformed with plasmid b365, expressing mCherry-Pex15(TA), and with plasmid pHS1, expressing Cox4pre-GFP. Cells were examined by fluorescence microscopy.

## REFERENCES

1. Surovtsev, I. V. & Jacobs-Wagner, C. Subcellular Organization: A Critical Feature of Bacterial Cell Replication. Cell 172, 1271–1293 (2018).

2. Hammer, S. K. & Avalos, J. L. Harnessing yeast organelles for metabolic engineering. Nat. Chem. Biol. 13, 823–832 (2017).

3. Schluter, A., Real-Chicharro, A., Gabaldón, T., Sanchez-Jimenez, F. & Pujol, A. PeroxisomeDB 2.0: an integrative view of the global peroxisomal metabolome. Nucleic Acids Res. 38, D800–5 (2010).

4. Gabaldón, T. Peroxisome diversity and evolution. Philos. Trans. R. Soc. Lond. B. Biol. Sci. 365, 765–73 (2010).

5. Islinger, M., Cardoso, M. J. R. & Schrader, M. Be different-The diversity of peroxisomes in the animal kingdom. Biochim. Biophys. Acta - Mol. Cell Res. 1803, 881–897 (2010).

6. Faust, P. L. & Kovacs, W. J. Cholesterol biosynthesis and ER stress in peroxisome deficiency. Biochimie 98, 75–85 (2014).

7. Dean, J. M. & Lodhi, I. J. Structural and functional roles of ether lipids. Protein Cell 9, 1–11 (2017).

8. Parsons, M. Glycosomes: Parasites and the divergence of peroxisomal purpose. Mol. Microbiol. 53, 717–724 (2004).

9. Kim, P. K. & Hettema, E. H. Multiple pathways for protein transport to peroxisomes. J. Mol. Biol. 427, 1176–1190 (2015).

10. Erdmann, R. Assembly, maintenance and dynamics of peroxisomes. Biochim. Biophys. Acta - Mol. Cell Res. 1863, 787–789 (2016).

11. Giannopoulou, E. A., Emmanouilidis, L., Sattler, M., Dodt, G. & Wilmanns, M. Towards the molecular mechanism of the integration of peroxisomal membrane proteins. Biochim. Biophys. Acta - Mol. Cell Res. 1863, 863–869 (2016).

12. Mayerhofer, P. U. Targeting and insertion of peroxisomal membrane proteins: ER trafficking versus direct delivery to peroxisomes. Biochim. Biophys. Acta - Mol. Cell Res. 1863, 870–880 (2016).

13. Schlüter, A. et al. The evolutionary origin of peroxisomes: An ER-peroxisome connection. Mol. Biol. Evol. 23, 838–845 (2006).

14. Gabaldón, T. et al. Origin and evolution of the peroxisomal proteome. Biol. Direct 1, 1–14 (2006).

15. de Duve, C. The origin of eukaryotes: a reappraisal. Nat. Rev. Genet. 8, 395–403 (2007).

16. Esser, C. et al. A genome phylogeny for mitochondria among α-proteobacteria and a predominantly eubacterial ancestry of yeast nuclear genes. Mol. Biol. Evol. 21, 1643–1660 (2004).

17. Nowack, E. C. M. & Weber, A. P. M. Genomics-Informed Insights into Endosymbiotic Organelle Evolution in Photosynthetic Eukaryotes. Annu. Rev. Plant Biol. 1–34 (2018). doi:10.1146/annurev-arplant-042817-040209

18. Embley, T. M. & Martin, W. Eukaryotic evolution, changes and challenges. Nature 440, 623–30 (2006).

19. Rivera, M. C. & Lake, J. A. The ring of life provides evidence for a genome fusion origin of eukaryotes. Nature 431, 152–5 (2004).

20. Gray, M. W. Mosaic nature of the mitochondrial proteome: Implications for the origin and evolution of mitochondria. Proc. Natl. Acad. Sci. 112, 10133–10138 (2015).

21. Roger, A. J. Reply to ‘Eukaryote lateral gene transfer is Lamarckian’. Nat. Ecol. Evol. 2018 (2018). doi:10.1038/s41559-018-0522-6

22. Husnik, F. & McCutcheon, J. P. Functional horizontal gene transfer from bacteria to eukaryotes. Nat. Rev. Microbiol. 16, 67–79 (2017).

23. Singer, A. et al. Massive Protein Import into the Early-Evolutionary-Stage Photosynthetic Organelle of the Amoeba Paulinella chromatophora. Curr. Biol. 27, 2763–2773.e5 (2017).

24. Lacroix, B. & Citovsky, V. Beyond Agrobacterium-Mediated Transformation: Horizontal Gene Transfer from Bacteria to. Eukaryotes. Curr. Top. Microbiol. Immunol. 351, 139–57 (2018).

25. Lutfullahoğlu-Bal, G., Keskin, A., Seferoğlu, A. B. & Dunn, C. D. Bacterial tail anchors can target to the mitochondrial outer membrane. Biol. Direct 12, 1–9 (2017).

26. DeLoache, W. C., Russ, Z. N. & Dueber, J. E. Towards repurposing the yeast peroxisome for compartmentalizing heterologous metabolic pathways. Nat. Commun. 7, 1–11 (2016).

27. Okreglak, V. & Walter, P. The conserved AAA-ATPase Msp1 confers organelle specificity to tail-anchored proteins. Proc. Natl. Acad. Sci. 111, 8019–8024 (2014).

28. Chen, Y.-C. et al. Msp1/ATAD1 maintains mitochondrial function by facilitating the degradation of mislocalized tail-anchored proteins. EMBO J. 33, 1548–1564 (2014).

29. Francisco, T. et al. Protein transport into peroxisomes: Knowns and unknowns. BioEssays 39, 1–8 (2017).

30. Elgersma, Y. et al. Overexpression of Pex15p, a phosphorylated peroxisomal integral membrane protein required for peroxisome assembly in S. cerevisiae, causes proliferation of the endoplasmic reticulum membrane. EMBO J. 16, 7326–7341 (1997).

31. Elgersma, Y., Van den Berg, M., Tabak, H. F. & Distel, B. An efficient positive selection procedure for the isolation of peroxisomal import and peroxisome assembly mutants of Saccharomyces cerevisiae. Genetics 135, 731–740 (1993).

32. Buentzel, J., Vilardi, F., Lotz-Havla, A., Gartner, J. & Thoms, S. Conserved targeting information in mammalian and fungal peroxisomal tail-anchored proteins. Sci. Rep. 5, 1–14 (2015).

33. Kuravi, K. Dynamin-related proteins Vps1p and Dnm1 p control peroxisome abundance in Saccharomyces cerevisiae. J. Cell Sci. 119 3994–4001 (2006).

34. Schuldiner, M. et al. The GET Complex Mediates Insertion of Tail-Anchored Proteins into the ER Membrane. Cell 134, 634–645 (2008).

35. Wróblewska, J. P. et al. Saccharomyces cerevisiae cells lacking Pex3 contain membrane vesicles that harbor a subset of peroxisomal membrane proteins. Biochim. Biophys. Acta - Mol. Cell Res. 1864, 1656–1667 (2017).

36. Agrawal, G., Fassas, S. N., Xia, Z. J. & Subramani, S. Distinct requirements for intra-ER sorting and budding of peroxisomal membrane proteins from the ER. J. Cell Biol. 212, 335–348 (2016).

37. Joshi, A. S. et al. A family of membrane-shaping proteins at ER subdomains regulates pre-peroxisomal vesicle biogenesis. J. Cell Biol. 215, 515–529 (2016).

38. Knoops, K. et al. Preperoxisomal vesicles can form in the absence of Pex3. J. Cell Biol. 204, 659–668 (2014).

39. Meinecke, M. et al. The peroxisomal importomer constitutes a large and highly dynamic pore. Nat. Cell Biol. 12, 273–277 (2010).

40. Montilla-Martinez, M. et al. Distinct Pores for Peroxisomal Import of PTS1 and PTS2 Proteins. Cell Rep. 13, 2126–34 (2015).

41. Cohen, Y. et al. The yeast P5 type ATPase, Spf1, regulates manganese transport into the endoplasmic reticulum. PLoS One 8, 1–14 (2013).

42. Lockshon, D., Surface, L. E., Kerr, E. O., Kaeberlein, M. & Kennedy, B. K. The sensitivity of yeast mutants to oleic acid implicates the peroxisome and other processes in membrane function. Genetics 175, 77–91 (2007).

43. Marelli, M. et al. Quantitative mass spectrometry reveals a role for the GTPase Rho1 p in actin organization on the peroxisome membrane. J. Cell Biol. 167, 1099–1112 (2004).

44. Kragt, A., Voorn-Brouwer, T., Van Den Berg, M. & Distel, B. Endoplasmic reticulum-directed Pex3p routes to peroxisomes and restores peroxisome formation in a Saccharomyces cerevisiae pex3Δ strain. J. Biol. Chem. 280, 34350–34357 (2005).

45. Tam, Y. Y. C., Fagarasanu, A., Fagarasanu, M. & Rachubinski, R. A. Pex3p initiates the formation of a preperoxisomal compartment from a subdomain of the endoplasmic reticulum in Saccharomyces cerevisiae. J. Biol. Chem. 280, 34933–34939 (2005).

46. Hoepfner, D., Schildknegt, D., Braakman, I., Philippsen, P. & Tabak, H. F. Contribution of the endoplasmic reticulum to peroxisome formation. Cell 122, 85–95 (2005).

47. van der Zand, A., Braakman, I. & Tabak, H. F. Peroxisomal membrane proteins insert into the endoplasmic reticulum. Mol. Biol. Cell 21, 2057–65 (2010).

48. Cohen, Y. et al. Peroxisomes are juxtaposed to strategic sites on mitochondria. Mol. BioSyst. 10, 1742–1748 (2014).

49. Veenhuis, M. & van der Klei, I. J. A critical reflection on the principles of peroxisome formation in yeast. Front. Physiol. 5, 110 (2014).

50. Delille, H. K. & Schrader, M. Targeting of hFis1 to peroxisomes is mediated by Pex19p. J. Biol. Chem. 283, 31107–31115 (2008).

51. Koch, A., Yoon, Y., Bonekamp, N. A., McNiven, M. A. & Schrader, M. A role for Fis1 in both mitochondrial and peroxisomal fission in mammalian cells. Mol. Biol. Cell 16, 507–786 (2005).

52. Gandre-Babbe, S. & van der Bliek, A. M. The Novel Tail-anchored Membrane Protein Mff Controls Mitochondrial and Peroxisomal Fission in Mammalian Cells. Mol. Biol. Cell 19, 2402–2412 (2008).

53. Chio, U. S., Cho, H. & Shan, S. Mechanisms of Tail-Anchored Membrane Protein Targeting and Insertion. Annu. Rev. Cell Dev. Biol. 33, 417–438 (2017).

54. Costello, J. L. et al. Predicting the targeting of tail-anchored proteins to subcellular compartments in mammalian cells. J. Cell Sci. 130, 1675–1687 (2017).

55. Kyte, J. & Doolittle, R. F. A simple method for displaying the hydropathic character of a protein. J. Mol. Biol. 157, 105–32 (1982).

56. Petersen, T. N., Brunak, S., von Heijne, G. & Nielsen, H. SignalP 4.0: discriminating signal peptides from transmembrane regions. Nat. Methods 8, 785–6 (2011).

57. Weir, N. R., Kamber, R. A., Martenson, J. S. & Denic, V. The AAA protein Msp1 mediates clearance of excess tail-anchored proteins from the peroxisomal membrane. Elife 6, 1–28 (2017).

58. Joshi, S., Agrawal, G. & Subramani, S. Phosphorylation-dependent Pex11p and Fis1p interaction regulates peroxisome division. Mol. Biol. Cell 23,1307–1315 (2012).

59. Herskowitz, I. Functional inactivation of genes by dominant negative mutations. Nature 329,219–222 (1987).

60. Meeks-Wagner, D. & Hartwell, L. H. Normal stoichiometry of histone dimer sets is necessary for high fidelity of mitotic chromosome transmission. Cell 44, 43–52 (1986).

61. Baker, A. & Schatz, G. Sequences from a prokaryotic genome or the mouse dihydrofolate reductase gene can restore the import of a truncated precursor protein into yeast mitochondria. Proc. Natl. Acad. Sci. 84, 3117–3121 (1987).

62. Lemire, B. D., Fankhauser, C., Baker, A. & Schatz, G. The mitochondrial targeting function of randomly generated peptide sequences correlates with predicted helical amphiphilicity. J. Biol. Chem. 264, 20206–20215 (1989).

63. Lucattini, R., Likić, V. A. & Lithgow, T. Bacterial Proteins Predisposed for Targeting to Mitochondria. Mol. Biol. Evol. 21, 652–658 (2004).

64. Hall, J., Hazlewood, G. P., Surani, M. A., Hirst, B. H. & Gilbert, H. J. Eukaryotic and prokaryotic signal peptides direct secretion of a bacterial endoglucanase by mammalian cells. J. Biol. Chem. 265, 19996–19999 (1990).

65. Walther, D. M., Papic, D., Bos, M. P., Tommassen, J. & Rapaport, D. Signals in bacterial beta-barrel proteins are functional in eukaryotic cells for targeting to and assembly in mitochondria. Proc. Natl. Acad. Sci. U. S. A. 106, 2531–2536 (2009).

66. von Heijne, G. Signal sequences The limits of variation. J. Mol. Biol. 184, 99–105 (1985).

67. Koonin, E. V. Horizontal gene transfer: essentiality and evolvability in prokaryotes, and roles in evolutionary transitions. F1000Research 5, 1805 (2016).

68. Huang, J. Horizontal gene transfer in eukaryotes: The weak-link model. BioEssays 35, 868–875 (2013).

69. Nowack, E. C. M. et al. Gene transfers from diverse bacteria compensate for reductive genome evolution in the chromatophore of Paulinella chromatophora. Proc. Natl. Acad. Sci. 113, 12214–12219 (2016).

70. Martin, W. Evolutionary origins of metabolic compartmentalization in eukaryotes. Philos. Trans. R. Soc. B Biol. Sci. 365, 847–855 (2010).

71. Schlötterer, C. Genes from scratch - the evolutionary fate of de novo genes. Trends Genet. 31, 215–219 (2015).

72. Kirschner, M. & Gerhart, J. Perspective Evolvability. Proc. Natl. Acad. Sci. U. S. A. 95, 8420–8427 (1998).

73. Keskin, A., Akdoğan, E. & Dunn, C. D. Evidence for amino acid snorkeling from a high-resolution, in vivo analysis of Fis1 tail-anchor insertion at the mitochondrial outer membrane. Genetics 205, 691–705 (2017).

74. Mozdy, A. D., McCaffery, J. M. & Shaw, J. M. Dnm1 p GTPase-mediated mitochondrial fission is a multi-step process requiring the novel integral membrane component Fis1p. J. Cell Biol. 151, 367–379 (2000).

75. Förtsch, J., Hummel, E., Krist, M. & Westermann, B. The myosin-related motor protein Myo2 is an essential mediator of bud-directed mitochondrial movement in yeast. J. Cell Biol. 194, 473–488 (2011).

76. Bateman, A. et al. UniProt: The universal protein knowledgebase. Nucleic Acids Res. 45, D158–D169 (2017).

77. Krogh, A., Larsson, B., Von Heijne, G. & Sonnhammer, E. L. L. Predicting transmembrane protein topology with a hidden Markov model: Application to complete genomes. J. Mol. Biol. 305, 567–580 (2001).

78. Ŏrs, Ş. T., Akdoğan, E. & Dunn, C. D. Mutation of the mitochondrial large ribosomal RNA can provide pentamidine resistance to Saccharomyces cerevisiae. Mitochondrion 18, 7–11 (2014).

79. Sikorski, R. S. & Hieter, P. A system of shuttle vectors and yeast host strains designed for efficient manipulation of DNA in Saccharomyces cerevisiae. Genetics 122, 19–27 (1989).

80. Prinz, W. A. et al. Mutants affecting the structure of the cortical endoplasmic reticulum in Saccharomyces cerevisiae. J. Cell Biol. 150, 461–474 (2000).

81. Ryan, K. R., Leung, R. S. & Jensen, R. E. Characterization of the mitochondrial inner membrane translocase complex: the Tim23p hydrophobic domain interacts with Tim17p but not with other Tim23p molecules. Mol. Cell. Biol. 18, 178–87 (1998).

82. Taxis, C. & Knop, M. System of centromeric, episomal, and integrative vectors based on drug resistance markers for Saccharomyces cerevisiae. Biotechniques 40, 73–8 (2006).

83. Lee, S., Lim, W. A. & Thorn, K. S. Improved Blue, Green, and Red Fluorescent Protein Tagging Vectors for S. cerevisiae. PLoS One 4–11 (2013).

